# Notes on the diet and size of *Cassiopea*

**DOI:** 10.1101/2025.01.18.633672

**Authors:** Kaden Muffett, Marta Mammone, Ramón D. Morejón-Arrojo, Andrea Toso, Lara M. Fumarola, Anabelle Marques Klovrza, Valentina Cardona, Serafin Mendez Geson, Claire E. Rowe, Anthony Bellantuono, Maria Pia Miglietta, Rachel Collin

## Abstract

Medusae of the genus *Cassiopea* are common components of tropical and subtropical coastlines globally. Despite the broad distribution of this benthic scyphozoan, much about their ecology remains poorly described. Here, we collected over 100 adult *Cassiopea* individuals from Panama, the eastern United States, Cuba, the Philippines, Italy and Australia to examine continuity and differences in their diet across space, and to investigate whether their unique lifestyle is reflected in their diet. We found the majority of prey items to be associated with the epibenthos. The recovered prey were supermajority crustaceans, mainly harpacticoid copepods, with pteropods, nematodes, and miscellaneous eggs common as secondary components. Within the gastrovascular cavity of a single medusa, we found up to 379 items. There was a limited relationship between medusa size and prey items. Location had an impact on gut content diversity and medusa size had a small impact on the number of taxa found within the gut. In some sites, prey were scarce or absent from all medusae sampled. Overall, we reaffirm the diet previously recorded for small medusae in Puerto Rico and show that similar components are common in large and small medusae from throughout the East and West Atlantic and the Philippines.

## 1. INTRODUCTION

*Cassiopea* spp., commonly known as the upside-down jellyfish, are present in near-shore benthic communities throughout the tropics and subtropics (Holland et al., 2004; Morandini et al., 2017). They are often distributed in large, long-lasting aggregations (called “smacks”) on silty sediment and near seagrass beds or artificial habitats such as shrimp farms (Thé et al., 2020). *Cassiopea* spp. are characterized by branching oral arms bearing secondary mouths and pigmented paddle-shaped appendages (Lampert, 2016) (Figure 1). Individuals can range from several centimeters to maximum sizes of 30 to 49 cm (Thé et al., 2020; Morejón-Arrojo & Rodríguez-Viera, 2023). Despite ubiquity in the tropics, *Cassiopea* are understudied in their natural habitat relative to laboratory specimens. These medusae rely on a mutualistic association with dinoflagellate microalgae (family Symbiodinaceae) for a large proportion of their energetic needs, providing the algae with a regular supply of inorganic molecules (carbon dioxide, ammonia) in return for carbohydrates and lipids of photosynthetic origin (Verde & McCloskey, 1998; Mortillaro et al., 2009). While microalgae are likely providing >100% of the *Cassiopea* carbon budget regardless of size, algal cell proliferation is supported by heterotrophy, thus *Cassiopea* requires a heterotrophic input (Verde & McCloskey, 1998; Stoner et al., 2016).

**Figure 1.**
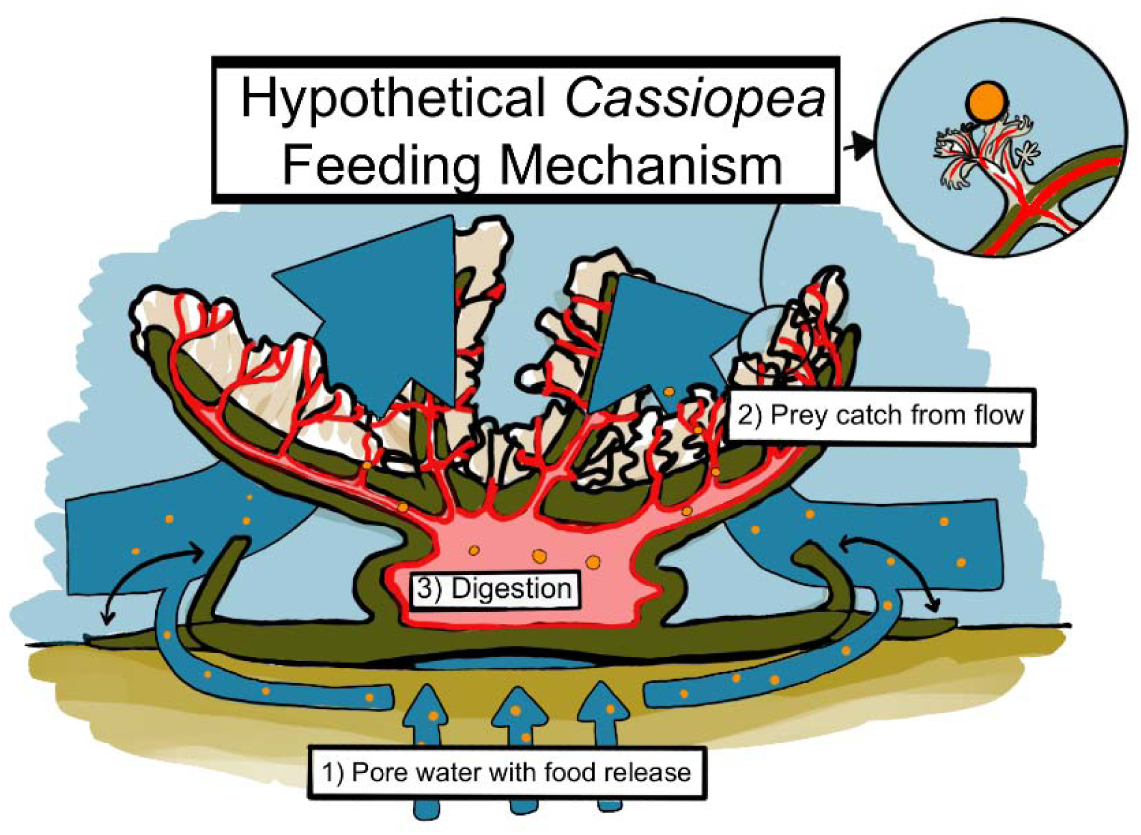
Diagram of water flow through a *Cassiopea* medusa as described in Durieux et al. 2023 and Santhanakrishnan et al. 2012. In order: 1) Pore water is siphoned from beneath the medusa during bell pulsation, then drawn over the bell margin and up through the oral arms, 2) Mouthlets immobilize prey with nematocysts, then draw prey into oral arms. 3) Food is digested in the gastrovascular cavity.

Due to their unique physiology, *Cassiopea* spp. represent hotspots of activity on the benthos, causing fluxes in oxygen and nutrient cycling through a combination of autotrophy, heterotrophy, and bioturbation (Welsh et al., 2009; Durieux et al., 2021). Of these factors, the heterotrophy of *Cassiopea* has been addressed in the literature only infrequently. *Cassiopea* spp. feed on zooplankton collected by secondary mouths when water currents flow above and through oral arms (Larson, 1997; Santhanakrishnan et al., 2012), as well as on POM (i.e., particulate organic carbon) and DOM (i.e., dissolved organic carbon) (Jantzen et al., 2010). *Cassiopea*’s unique morphology likely provides access to a unique feeding strategy, but the details of this strategy remain largely hypothetical (Figure 1) (Dureiux et al. 2023). As temperature and light vary across time and space, *Cassiopea* may shift from autotrophy to heterotrophy and vice versa, allowing *Cassiopea* spp. to adapt and adjust to conditions. For example, efficient heterotrophic feeding allows *Cassiopea* to mitigate decreased photosynthate translocation during heat stress (Djeghri et al., 2021; Muffett et al., 2022; Thé et al., 2023; Toullec et al., 2024).

Historically, the trophic role of scyphomedusae has been oversimplified (Pauly et al., 2009; Nagata et al., 2024) and minimized over time, with an emphasis historically on only pelagic *Aurelia* spp. And *Chrysaora* spp. and more recently on the Mediterranean barrel jellyfish *Rhizostoma pulmo* (Leoni et al., 2022).

Here, we present dietary information of the benthic jellyfish *Cassiopea* spp. collected from the waters of Florida (Key Largo), Cuba (Havana), Panama (Bocas del Toro), Philippines (Bolinao), and Italy (Sicily). With additional notes on three collections with limited gut contents (one in Australia, two in Florida) and information on *Cassiopea* populations around the world with data on their size, weight, and sex distribution.

## 2. METHODS

### 2.1 *In situ* Gut Contents Collections

We analyzed the gut content of 130 medusae collected from eight locations (Table 1 and Figure 2).

**Figure 2.**
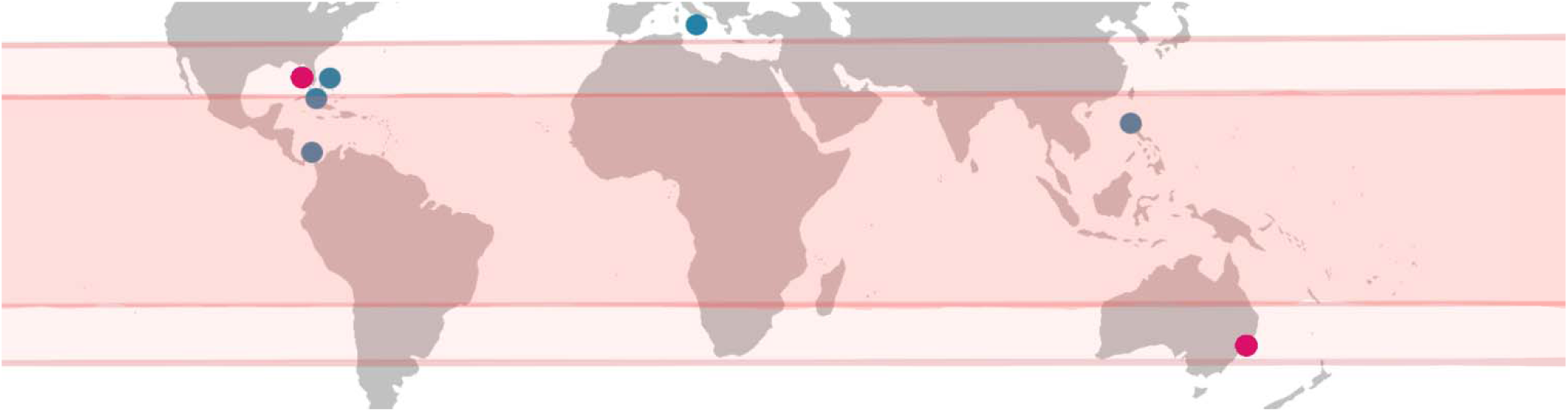
Global map highlighting collection points across the world. Demarcations for the tropics and subtropical latitude boundaries are listed. High-quality diet collections (blue) and low-quality (minimal prey) diet collections (pink) are displayed.

**Table 1.**
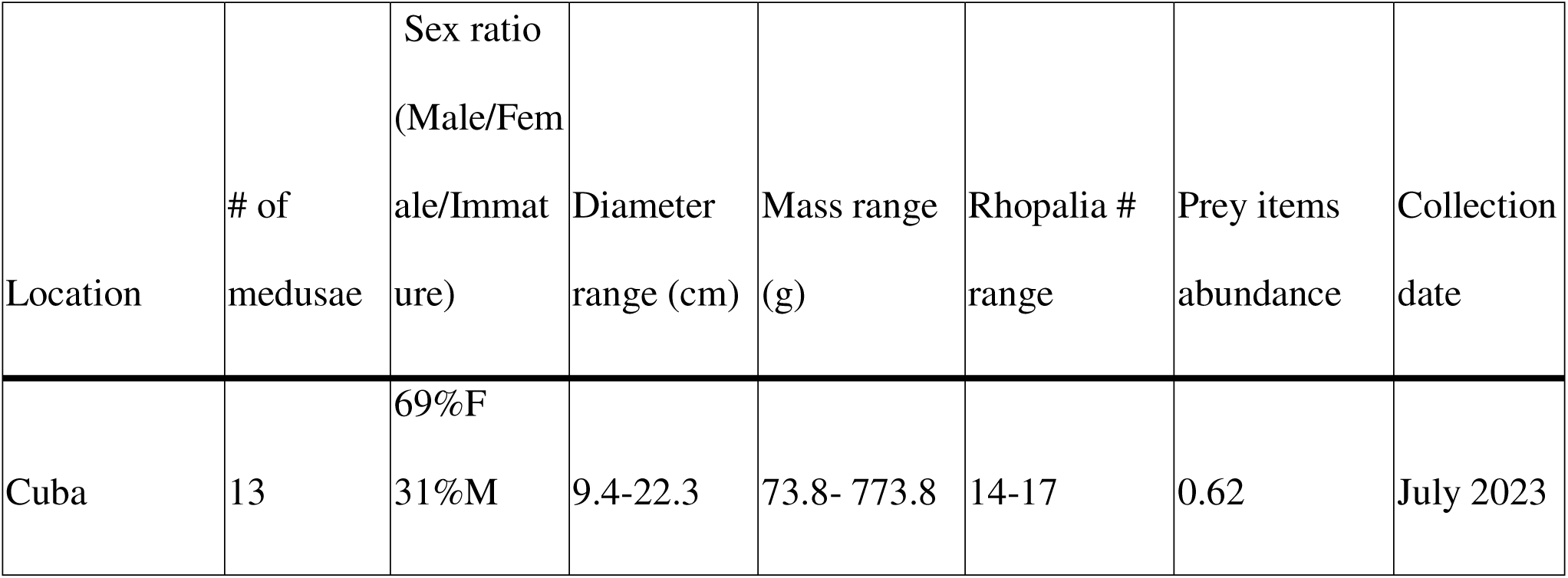
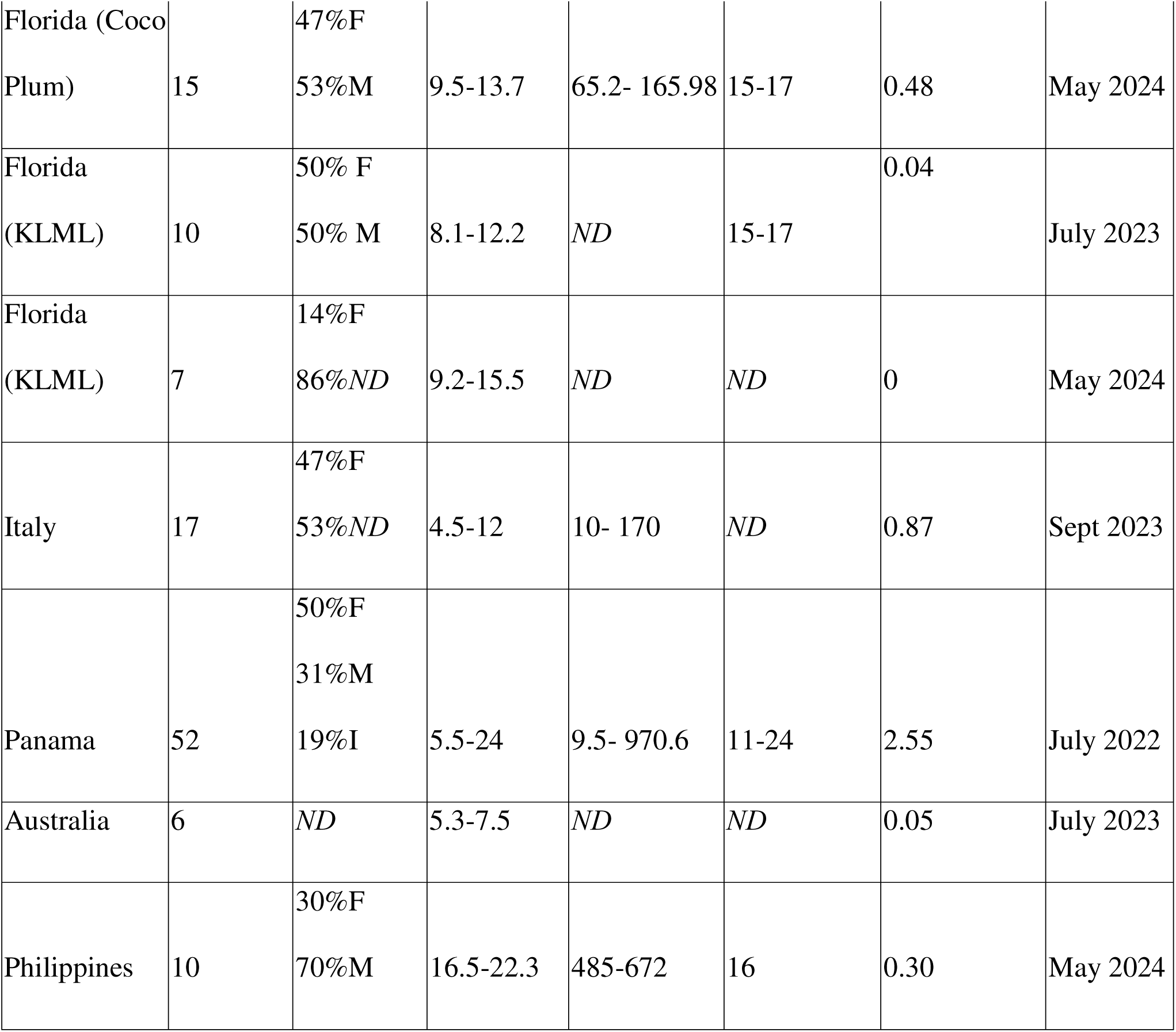
Details of collections including location, number of medusae, sex of collected medusae expressed as percentages, range of diameters for medusae, mass in grams, number of rhopalia if determined and prey abundance. Prey abundance is scaled to average medusa size at location (average total items/average bell diameter). Missing information is listed as ND.

Fifty-two medusae (*Cassiopea xamachana*) were collected from three sites within the seagrass beds offshore of the Smithsonian Tropical Research Institute, Bocas del Toro, Panama at 1 to 2 m depth (Lat: 9.3513 to 9.3523, Long: -82.2571 to -82.2580) in July 2022 during daylight hours. Medusae were collected from three specific sites within this study area, a high-light sandy site near a dock, a high-light sandy site adjacent to a seagrass patch, and a lower-light silty site along the mangrove edge.

Fifteen medusae (*Cassiopea xamachana*) were collected from a silty, shallow, calm inlet on Coco Plum Rd. in Key Largo, Florida, US (∼1m depth) (Lat: 27.079, Long: -80.452). Collections occurred in May 2024 during the early afternoon.

Thirteen medusae (*Cassiopea* cf. *xamachana*) were collected from a shallow sandy mangrove stand outside of Havana, Cuba (∼ 0.4-0.8 m depth) (Lat: 23.068, Long: -82.528) during morning hours in July 2023.

Seventeen medusae (*Cassiopea andromeda*) were collected from the Palermo port at 0.5 m depth in anoxic muddy sediment near a waste outflow (Lat: 38.11966, Long: 13.36691). Gut content was analyzed for 11 out of 17 and only weight and measurements were taken for the remaining six. Collections occurred in September 2023 during afternoon hours (seawater temperature of 25°C).

Ten medusae (*Cassiopea* cf. *xamachana*) were collected from silty seagrass beds at Key Largo Marine Laboratory (Lat: 25.101141, Long: -80.43842) during afternoon hours in July 2023 during a marine heat event (36 °C) at a depth of greater than 1m. Medusae were visibly bleached and in active decline. Medusa gut content was fixed in formalin and stored until analysis (Jun 2024). In May 2024, seven additional medusae were collected from this site in the morning hours and dissected for gut contents during less extreme temperature conditions (31°C).

Ten medusae (*Cassiopea* sp.) were collected from a sheltered, nearshore (∼ 2m depth) silty-sandy habitat in the Guiguiwanen Channel, Bolinao, Philippines (Lat: 16.383482, Long: 119.923245) during morning hours in May 2024. This area is heavily impacted by mariculture fish cages, leading to eutrophic conditions. Gut content was fixed in formalin until analysis (June 2024).

Six medusae (*Cassiopea* sp.) were collected from Lake Macquarie, New South Wales, Australia (Lat: - 33.123094, Long: 151.538484) during the early afternoon in July 2023. Medusae were collected from muddy sediment at 30-45 cm depth in 18 °C temperatures. Medusae were small and collected at the end of their season.

In Panamanian, Cuban and Florida Keys collections, oral arm morphology, oral arms anatomy, shape of the vesicles indicated that collected samples were not *C. frondosa* (Fitt et al. 2021; Larson, 1997; Morandini et al., 2017). In the Florida Keys, different species of *Cassiopea* coexist (Fitt et al. 2021; Muffett et al., 2023). Thus, a subset of samples were fixed in ethanol for barcoding.

Where possible, medusae were photographed *in situ,* hand collected in a bucket and weighed; the diameter of each specimen was measured with a ruler and rhopaliar number was also recorded (see supplementary table). In addition, information such as time of day of collection, time of collection, and location characteristics were collected. To analyze the gut content, an incision across the oral arms was performed to access the stomach cavity, and stomach contents were siphoned out with a syringe or pipette, categorized under major taxonomic groups and counted under a standard 50x magnification scope.

To understand the approximative degree of contamination caused by laboratory supplies such as plastic pipettes, in Panama, some water blanks were made by taking three 5 mL aliquots of the sink water, agitating it with the plastic pipette that was used for gut content and examining it under the scope counting visible plastics.

Female specimens of *Cassiopea* spp. are usually recognizable by the presence of a white oral appendages cluster in the center of oral arms (Figure S1), however, for Panama and Florida (Coco Plum) samples, sex was confirmed by microscopic identification of gonadal tissue sampled with a tweezer after siphoning gut content.

### 2.3 Barcoding

DNA was extracted from four samples each from the Panama and Florida Keys collection to determine the *Cassiopea* species. Extractions were done following the protocol in Frolova et al. 2022 for Floridian (Coco Plum collection) samples and using Zymo QuickDNA kit for Panamanian samples.

PCR was run using BioRad T100 thermal cycler (2min 94°C, [30s 94°C, 30s 48°C, 1min 72°C]X34, 7min 72°C) using primers Cass 12037 (5’-ATYAGGAGCAGGATTCAGTATG-3’) and Cass 12584 (5’-CTGCTGGGTCAAAGAATGAGG-3’). After amplification was confirmed with a gel, PCR products were diluted 1:10 with nuclease-free water and used for the Cycle sequencing PCR (1min 95°C, [[10s 95°C, 5s 50°C, 1:15min 60°C]X14, [10s 95°C, 5s 50°C, 1:30min 60°C]X4, [10s 95°C, 5s 50°C, 2min 60°C]X4]). Cycle sequencing products were then purified with Zymo ZR DNA sequencing Clean-up Kit before running on seqStudio Genetic Analyzer (Thermo Fisher Scientific). Species identities could not be confirmed for Cuban, Australian, Italian, and Phillipines collections.

### 2.4 Data analysis

Benthos and zooplankton assignments were made based on WoRMS attributes of lowest identified taxon (zooplankton, benthic or NA/mixed).

Foraminifera, detritus pellets and microplastics were excluded from counts of total prey items as likely accidental uptake.

Data analysis was performed in R v. 4.2.2 (http://www.R-project.org/) using RStudio v. 2023.12.1. The package “vegan” was used to compute Simpson and Shannon diversity indices. Linear regression and ANOVA were run with the “stats”, then selected with the “MuMin”, “rcompanion”, “ggfortify” and “car” packages(Mangiafico, Tang et al. 2016, Fox & Weisberg 2019, R-Core-Team (Foundation for Statistical Computing) 2022, Bartoń 2024). Total prey were log-transformed using the log function for normality. For values that were not homoscedastic and normal across sites base on testing (Levene Test (leveneTest) and Shapiro test (Shapiro.test)), Kruskal Wallis tests with Dunn comparisons were performed to compare medians (“FSA” package)(Ogle et al. 2023). and Graphs were built using the “ggplot2”, “plyr”, “ggstatsplot”, “cowplot” and “ggpubr” packages (Wickham 2016, Patil 2021, Kassambara 2023). Best fit lines of weight versus diameter were compared using the trendline functions in Google Sheets. Dietary barplots were made in Excel. Project code is available on Github (https://github.com/kadenmuffett/Cassiopea_gut_content).

## 3. RESULTS

### 3.2 Gut content diversity

An average of 19 (median: 9, range: 0-379) prey items were collected from each medusa examined. Medusa diameter had a significant but limited impact on total number of prey recovered (Pearson’s coefficient correlation, R=0.283, t = 2.93, df = 99, p-value = 0.004) (Figure 3). A total of 1921 prey items were examined, of which the vast majority (75.8%) were crustaceans. As crustacean identification requires specialized expertise and many crustaceans were partially digested, we report crustaceans in generalized categories (e.g. harpacticoid copepods, cyclopoid copepods, mystacocarids) based on observed body plan (IMOS Zooplankton Identification Guide).

**Figure 3.**
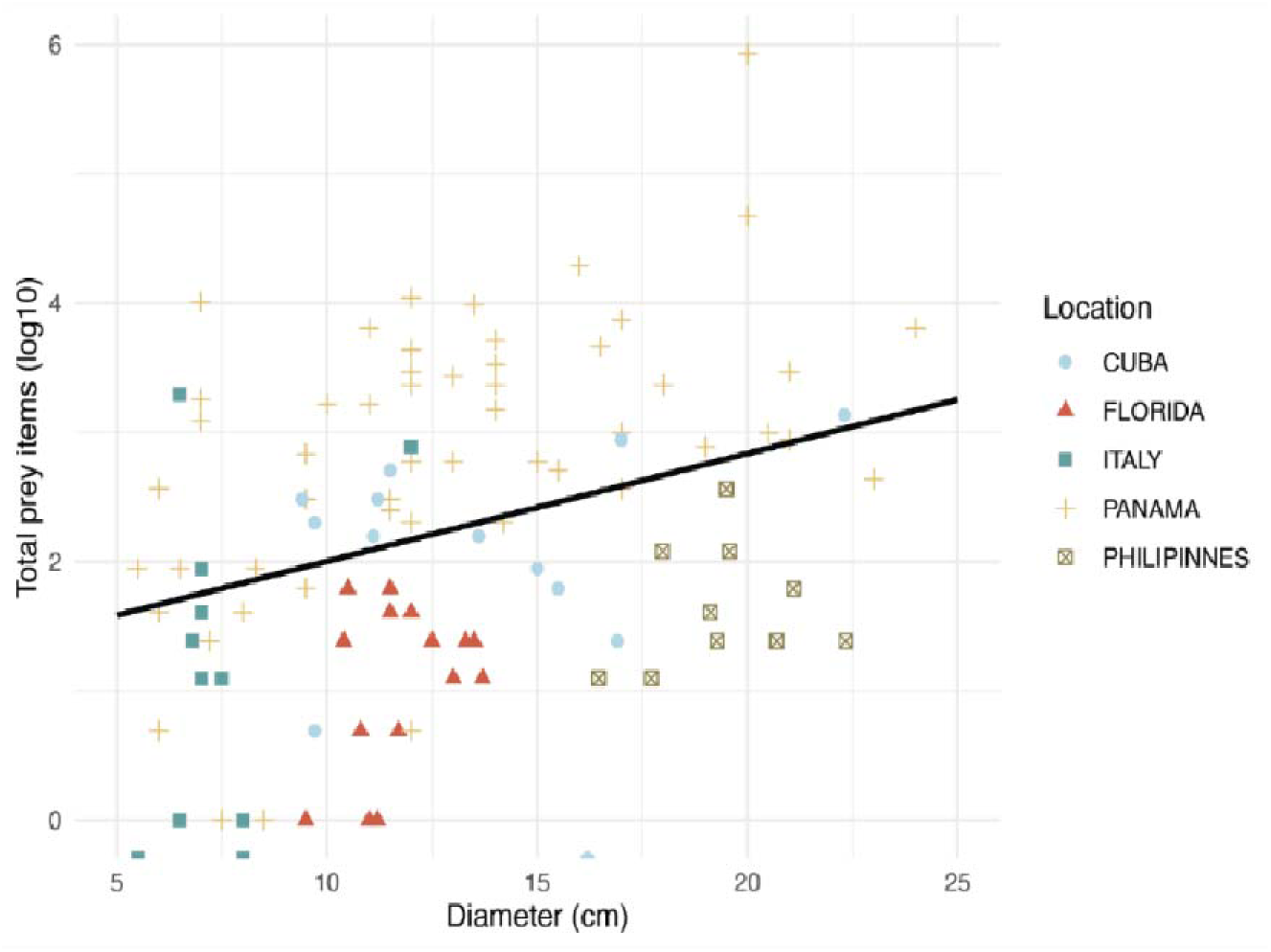
*Cassiopea* and prey recovered by size of medusa, shows a slight upward trend in prey by diameter (R = 0.28, p = 0.004). Size and color represent collection location. Extremely low yield collections are not included (Florida KLML and Australia).

In addition to crustaceans, nematodes, molluscs, pteropods, and eggs (primarily those resembling copepod egg masses, but also larger miscellaneous eggs) were found in medusa gastrovascular cavities (Fig 4 and S2). Foraminiferans were present in 24 individuals in Cuba and Panama. Insects were found in the Cuban and Floridian medusae, potentially related to the proximity to mangroves and the shallowness of the site where collections occurred.

**Figure 4.**
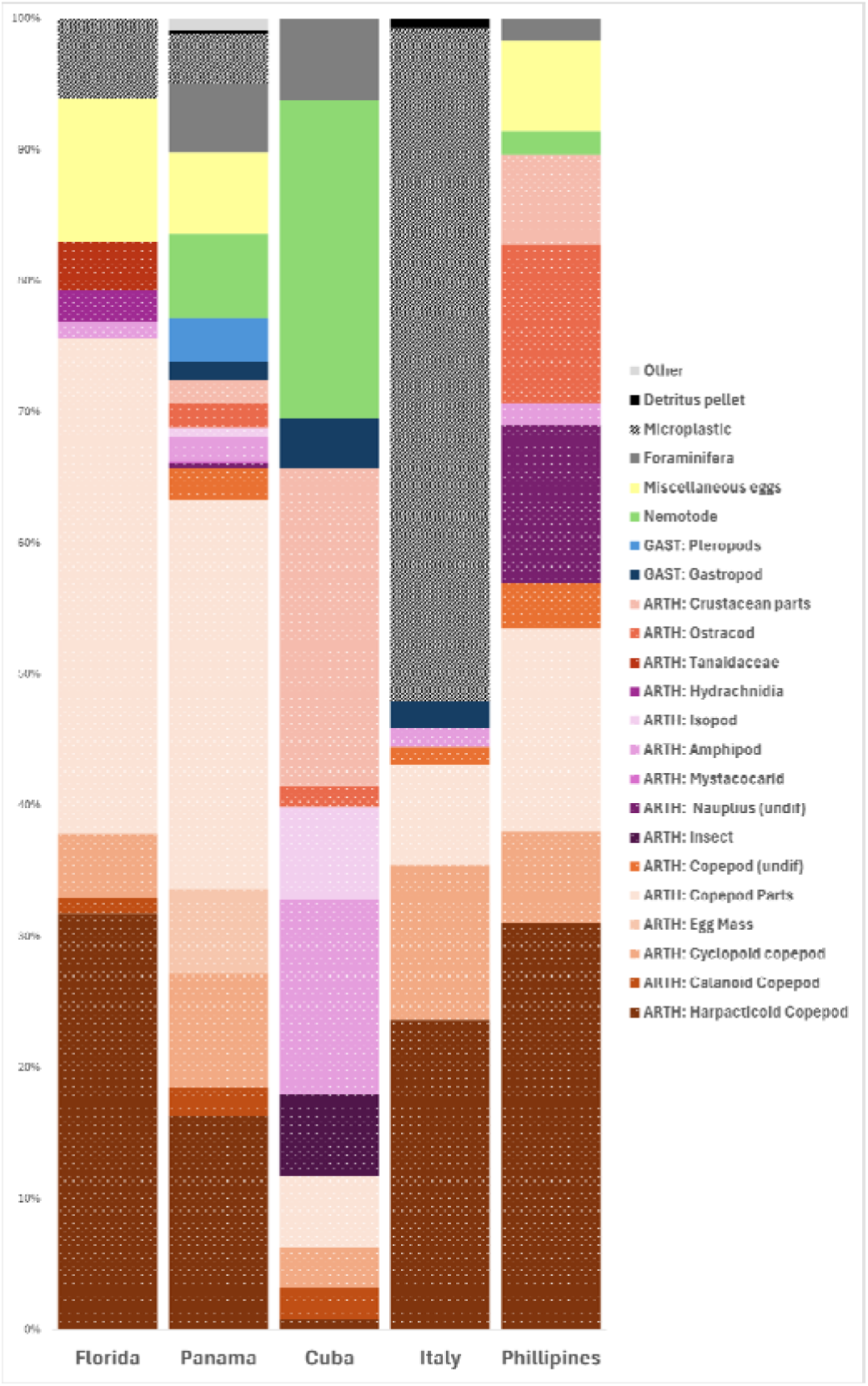
Stacked bar plot of items found in gastrovascular cavities aggregated by medusa location. Dotted columns are arthropods.

Microplastics were also found within the gastrovascular cavities of medusae but were not more prevalent than those found in water blanks generated in the laboratory for Panama samples. Microplastics were most abundant within GVCs of medusae from the Italy dock site, where they represented 52% of the retrieved content (see Figure 4).

The Shannon diversity varied across the locations tested (Kruskal-Wallis test, *χ*^2^= 31.12, p=2.89e-06). The mean Simpson diversity was 0.635 +/- 0.249. The mean Shannon diversity was 1.150 +/- 0.638 and was higher in Cuba, and Panama samples than in the Floridian and Italian samples (see KW Holm p-vals in Figure 5).

**Figure 5.**
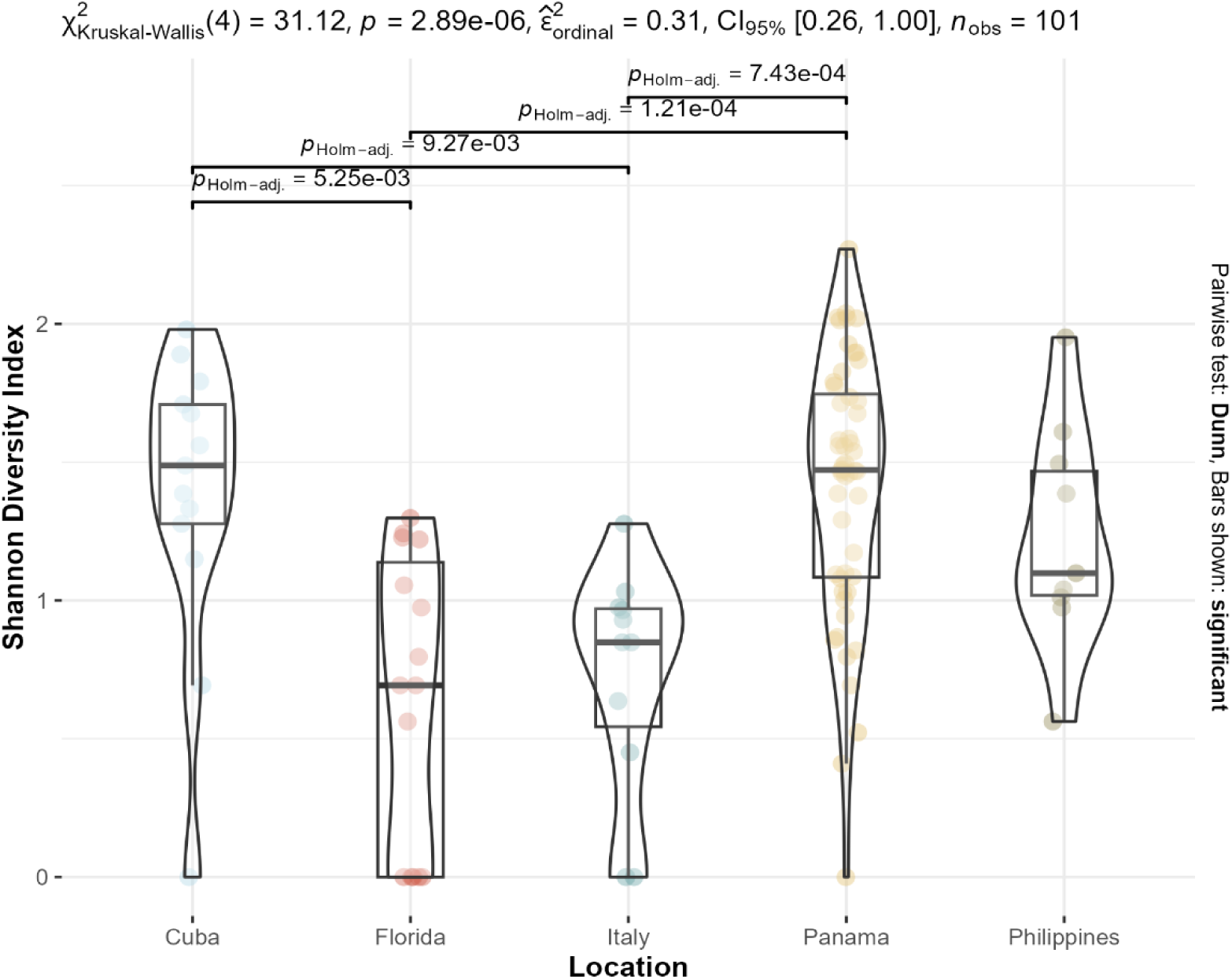
Shannon diversity by site location. Mean is represented by a center line. Significant pairwise comparisons of the Kruskal-Wallis test are presented with lines above violin plots.

Diversity was similar across site types (muddy, sandy, or silt/sand combination) after accounting for location (ANOVA type II, df=1, f=1.98, p=0.16) (Figure 6).

**Figure 6.**
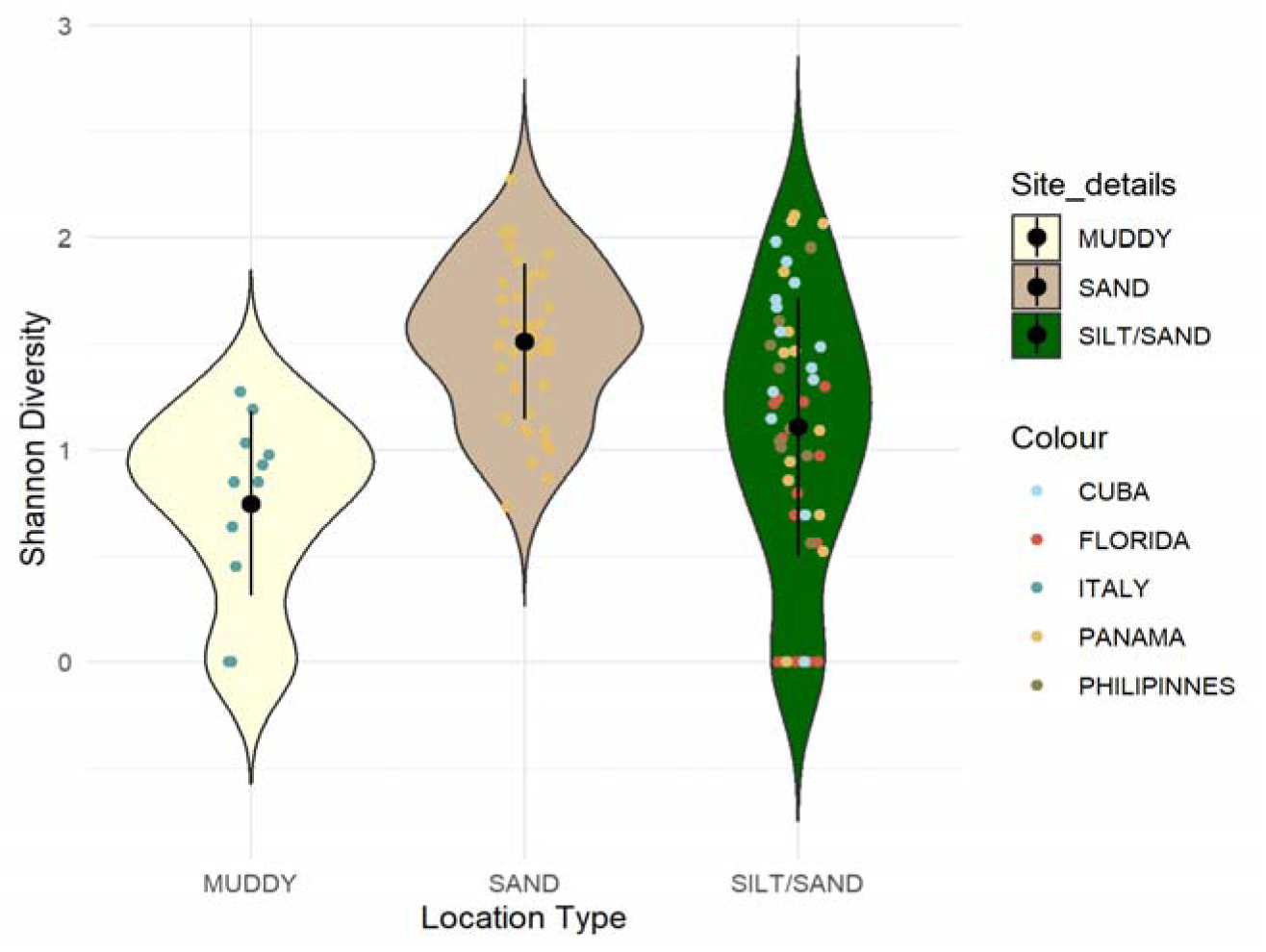
Shannon diversity by location type. Dot color indicates location while violin plot color indicates site type. The black center represents the mean with one standard deviation in either direction represented by the black center line.

### 3.2 Gut content members

Specific dietary components varied by location, harpacticoid copepods were highest within Italian samples and lowest in Cuban samples. Floridian, Panamanian, Italian and Philippines samples did not have significantly different proportions. Arthropods overall were significantly reduced in Cuba, with the median proportion of the recovered diet in Italy, Florida and the Philippines 100% arthropods. Copepods alone had a median of 71% or greater in Panama, Florida and Italy (Fig 7).

**Figure 7.**
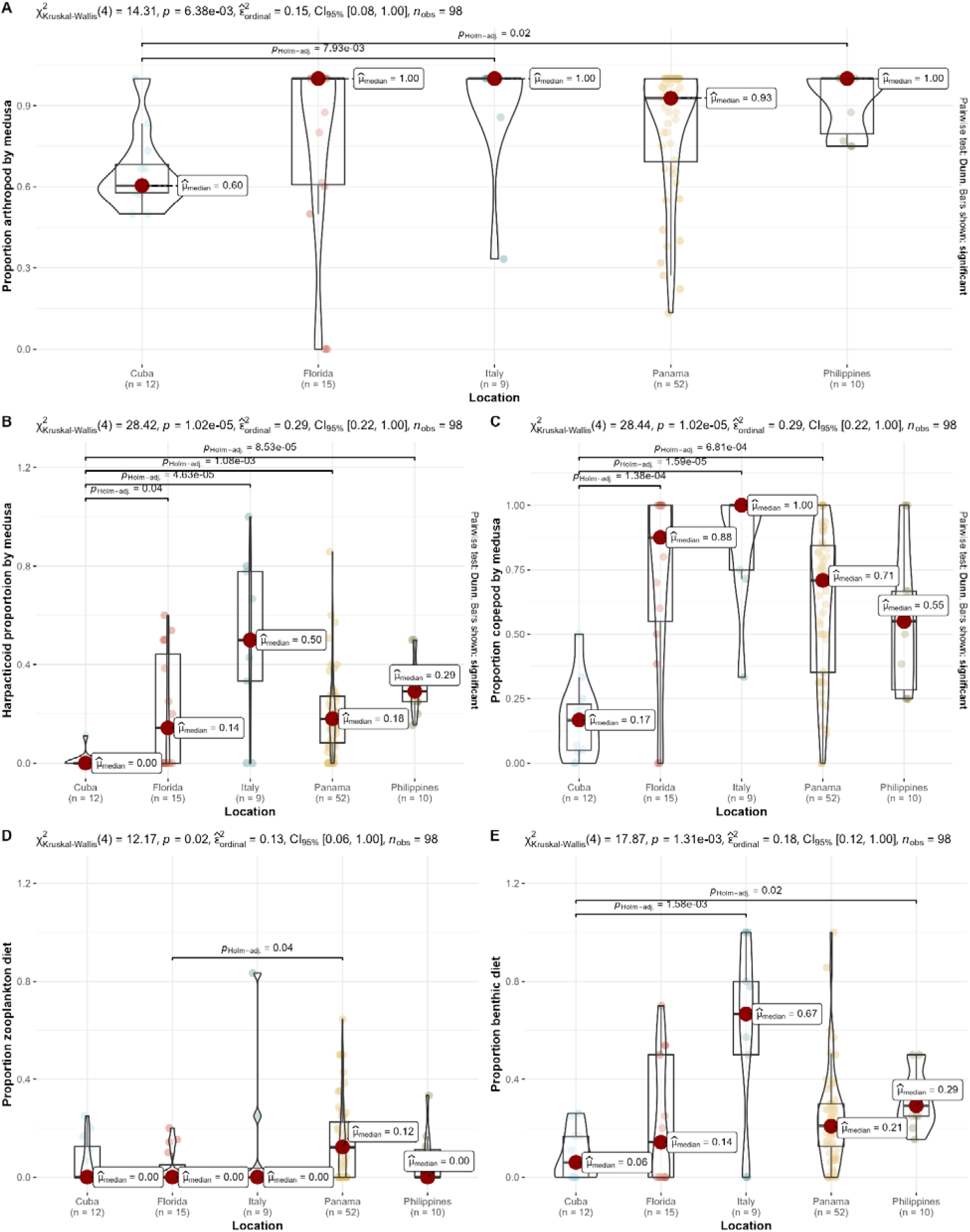
Comparisons between location of collection and proportion of food collected for (A) overall proportion arthropod abundance, (B) proportion of harpacticoids, and (C) proportion copepods. (D) Zooplankton and (E) Benthic represent proportion of the identified prey that could be assigned to these groups according to WoRMS as of Dec 2024. All test statistics shown are Dunn test pairwise comparisons with holm correction. Nonsignificant p-values (>0.05) are not displayed.

A mean of 26% of *Cassiopea* dietary contents could be assigned as meiobenthos or benthos. A mean of 11% were assigned as zooplankton. Cuba had a slightly lower proportion of benthos-associated taxa relative to the Philippines and Italy (KW chi2, Holm adj Dunn, Phil. p=0.02, It. p=0.0016).

### 3.3 Low prey collections

In both Australia and Florida (KLML), medusae had very little or no prey items in their GVCs (see Table 1). The six medusae from Australia were collected in winter and were quite small (diameter range: 5.3-7.5cm) (Figure 8). Only two total copepods were collected from their GVCs (prey abundance =0.30, see Table 1).

**Figure 8.**
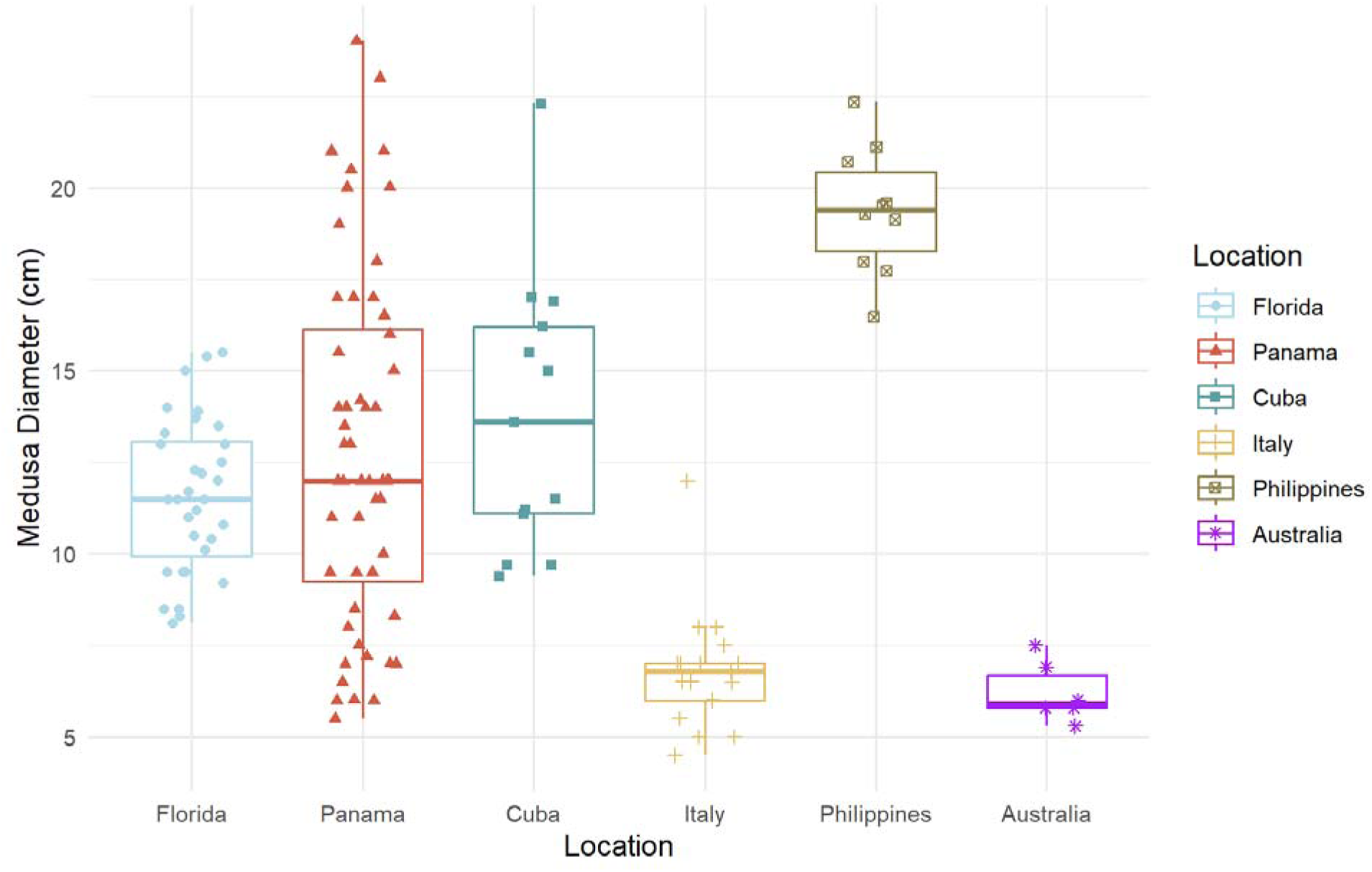
Diameter of Cassiopea by location of collection. The Florida graphed box includes all the Florida locations.

In Florida, animals were collected at the Key Largo Marine Research Laboratory in July 2023, during the 2023 Florida Keys marine heat event, and in May 2024. Four prey items were retrieved from the 2023 collection and none from the 2024 collection (prey items abundance = 0.04 and 0 respectively, see table 1). In July 2023, one insect, one tanaid, and two detritus pellets were found across 10 total medusae.

### 3.1 Summary of medusa metrics

Among individuals collected in Panama, immature medusae ranged from 5.5 to 8.3 cm in diameter, definite females from 14 to 24 cm in diameter, and males from 11 to 21 cm. While medusae smaller than 5.5 cm were present, they were excluded from diet analysis as the interior of their gastrovascular cavities (GVCs) proved too difficult to probe. Overall (mean diameter), medusae from Philippines were larger than those collected in the other sampling efforts (Figure 8).

Medusa’s weight increased with diameter. Weight (g) increases were best explained by a power series of diameter (D) across all medusae (g = 0.111*D^2.83^, R^2^=0.951) (Figure 9). When tested against extremely small ephyrae (dataset from Muffett 2021), this model produces a small overestimate in weight within this size class. The graph of the revised power series including data at both ends of the *Cassiopea* size scale (g=0.0557D^3.1^, R^2^= 0.993) is available in the supplementary materials (S3).

**Figure 9.**
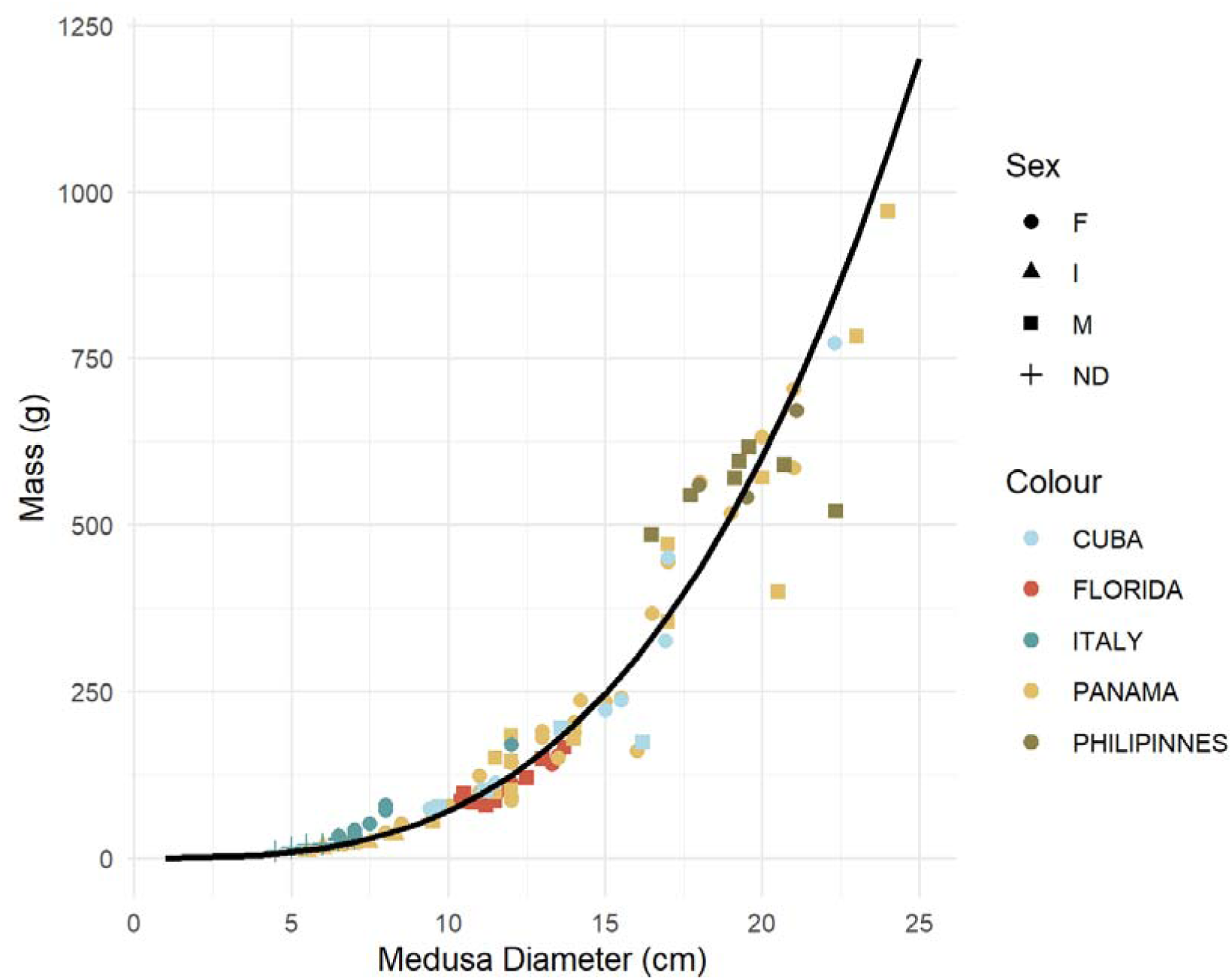
Diameter and weight of medusae with location (color) and sex (shape) details. Sites and individuals with no mass data are excluded. F=female; M=Male; I=immature; ND=not determined.

While observed sex distribution varied between locations, not all locations verified gonadal maturity.

## 4. DISCUSSION

### 4.1 Gut Content

Our results show that the diet of *Cassiopea* is heavily skewed towards epibenthic fauna (harpacticoid amphipods, nematodes, foraminifera), reinforcing the current understanding of these medusae as competent sediment feeders. While regionally limited dietary work on pelagic scyphozoans has shown a high prevalence of pelagic copepods as food sources, our results show that *Cassiopea*’s benthic behavior is reflected in its diet characterized by copepod groups that are a key dietary resource (Larson 1997; Graham & Kroutil, 2001; Nagata & Morandini, 2018; Nagata et al., 2024).

Overall, the majority of recoverable gut contents of *Cassiopea* spp. are crustaceans, a group dominated by copepods at all sites other than Cuba. In Larson (1997), *Cassiopea* from La Parguera (Puerto Rico) were reported to feed upon copepods (mostly harpacticoids) and ostracods in highest abundance with supplemental tanaids and chironomids. Other prey consisted of nematodes, mysids, miscellaneous eggs, pteropods, chaetognaths, cumaceans and foraminifera. Inconsistent with Larson were the insects found in *Cassiopea* specimens from Cuba and Florida (during 36 °C collection). While rare, other medusae have been reported with terrestrial food items elsewhere (e.g. the hydromedusae *Maeotias marginata* (Modeer, 1791) see (Wintzer et al., 2011). The presence of insects in *Cassiopea* from Cuba might be related to the proximity of the populations to the mangrove roots, as mangroves are considered a hotspot of insect diversity (Yeo et al., 2021).

The total number of prey was also explained by the jellyfish diameter (p=0.004), with bigger jellyfish containing higher amount of preys in their gut. This is in discordance with previous studies on *C. xamachana* from the Bahamas that shows that neither dry weight of food in the gut nor microalgal endosymbionts density are correlated to bell diameter (Stoner et al., 2016).

Diversity indices of gut content varied significantly by site, though substrate type was not a primary driver. Specific gut components varied with site. Cuba had a lower number of harpacticoid copepods but had higher prevalence of nematodes and amphipods. Italian samples had a high proportion of inorganic plastic pollution within their bodies. The Guiguiwanen Channel (Philippines) has been characterized as highly eutrophic, primarily due to the significant organic load from excessive fish feed inputs associated with mariculture activities. This has resulted in notable shifts in the phytoplankton community and frequent algal blooms (San Diego-McGlone et al., 2008). While the broader ecological significance of this remains elusive, these eutrophic conditions may be linked to the higher Shannon diversity observed in the Philippine *Cassiopea* samples, possibly providing a more diverse range of prey items. Additionally, the elevated nutrient levels from eutrophication have been correlated with increase in *Cassiopea* sp. medusae biomass (Stoner et al., 2022). However, water quality conditions in Cuba and Panama, which might influence similar diversity patterns, remain uncertain.

Gut content isolation often (3/8 locations) recovered little prey. Whether this represents true ecological states, or a quirk of sampling requires more sampling attempts to be properly assessed.

### 4.2 Estimating overall consumption

*Cassiopea* is characterized by a unique lifestyle, living upside-down which brings to a unique feeding behavior that involves benthic nutrient fluxes. Rhizostomeae, like *Cassiopea*, have branching oral arms with secondary mouths. *Rhizostoma pulmo*, a pelagic rhizostome, was shown to have 253 ±338 prey items per adult medusae (Leoni et al., 2022). Our data showed an average of 19 prey items per specimen (Table 2). This distinction could be attributed to the differing lifestyles (benthic vs pelagic) and sizes of the medusae examined in both investigations. In addition, medusae may change their diet in different ways throughout their ontogeny (Nagata et al., 2024). Unlike *Cassiopea*, *R. pulmo* gut content was characterized by micro and meso-zooplankton, with ciliates and calanoid copepods dominating respectively (Leoni et al., 2022).

**Table 2.** Description of total sampled medusae (n=130) including mean and range for diameter, mass, and total prey items/medusa, as well as the proportion of male/female medusae, as sexed by gonad observation. Not all medusae were sampled for all measurements.

| Parameter | Mean (+/- sd) or % in Sample Population |
| --- | --- |
| Diameter (cm) | 12.04 (+/- 4.72) |
| Mass (g) | 208.8 (+/- 220.3) |
| Sex | Male (32.7%), Female (49.5%), Immature (9.3%), Undetermined (8.4%) |
| Total prey | 19 (+/-40) |

For medusae of the genus *Stomolophus*, active planktonic filter feeders, the closest relative of *Cassiopea* for which we have comparative data, prey digestion time is approximately 1.5-2 hours (Arai, 1997). As such, our collections likely represent a maximum of 2 hours of prey capture by *Cassiopea*. Moreover, our gut content analysis did not recover prey that might have been attached to oral arms or within the oral arm canals. Assuming consumption occurs for approximately 12 hours per day with the assumption that overnight quiescence results in limited consumption (Nath et al., 2017), and our mean of 19 prey items, a medusa may consume roughly 114 prey items daily. This is very likely an underestimate.

Gut residence times of the different dietary components vary, thus our ability to draw generalizations is limited. Both within narrow and broad temperature ranges, scyphozoan digestion speeds up with temperature (Arai, 1997). Overall high digestion rates may negatively impact gut content recovery even if prey number is similar.

### 4.3 Nonnative species

The non-native amphipod *Laticorophium baconi* (Shoemaker, 1934) was identified within Italian samples. *L. baconi* has been reported as introduced globally in Hawaii, the Eastern US, the Gulf of Mexico, China, Brazil, New Zealand, and Australia (Barnard, 1970; LeCroy, 2004; Hirayama, 1990; McKinney, 1977; Storey, 1996; Sylvester et al., 2011; Valério-Berardo & de Souza, 2009). In 2018, it was found for the first time in European waters, spreading since, to Mediterranean countries including Italy. This amphipod inhabits fouling communities on vessels but also within port environments, artificial structures, and human-impacted areas (Guerra-Garcia et al., 2023) As *Cassiopea andromeda* is a similarly recent entrant to the Italian coast (Cillari et al., 2018), their cooccurrence reinforces the alteration of the Mediterranean food web that remains ongoing.

### 4.4 Low prey collections

In both Australia and Florida (KLML), medusae had little prey. As the Australian medusae were collected in winter, medusa feeding may have been altered by low temperatures. These data do indicate that a small amount of feeding continues into the winter, even if medusa health fails. In fact, as winter approaches and temperature drops, bell pulsation decreases impacting *Cassiopea*’s ability to feed (Rowe et al., 2022).

In Florida, the 2023 collection was during a regional coral bleaching event. While the low food abundance during the bleaching event of 2023 may indicate stress, a failure also to recover diet items from medusae in 2024 suggests that additional factors may be controlling feeding in this region. Medusae in heat stress had few or no food items, demonstrating that the temperature may reduce heterotrophic capacity. In the 2024 samples, the absent prey may be explained by algal overgrowth. The medusae were collected during a benthic algal bloom in the area. Of the medusae collected, six were not in direct contact with the sediment. In cases where extreme weather (heat) is followed by overgrowth, medusae may have especially high difficulty reestablishing and feeding without adequate substrate.

### 4.5 Laboratory diets

This work bears comparison to currently accepted laboratory conditions. When kept in the laboratory, *Cassiopea* spp. are often fed with *Artemia franciscana* or *Artemia salina* nauplii, the larval stage of a crustacean species commonly used in aquaculture (Lavens & Sorgeloos, 2000). These nauplii decrease in nutritional quality with days post-hatch and may fall short in matching adult medusa nutritional needs. While *Artemia* spp. seems to improve growth relative to other food types (e.g. *Spirulina maxima*, a slurry of *Mytilus galloprovincialis*, or polychaete-based experimental diets) (Mammone et al., under review), mixed foodstuffs or supplementation with larger prey items may be warranted when attempting to culture larger medusae, coupled with light that sustains photosynthesis.

### 4.6 Future directions

As discussed above, our analyses identify gastrovascular cavity contents but fail to differentiate between digestible and indigestible fractions. For future work, gut content examinations should be bolstered with stable isotope and fatty acid analyses, to assess the relative contribution of epibenthic and meiofauna to *Cassiopea* tissues during their ontogeny. In addition, prey should be counted also from oral arms and within gastrovascular canals.

Within the examined gastrovascular cavities, microfibers were identified in high abundance in Italian samples. Microplastics including plastic that are < 5 mm in length are increasingly found in nearshore systems (Auta et al., 2017). Jellyfish can ingest microplastics that can be present in the water column or sink to the bottom such as polyvinyl chloride (PVC), nylons, and polyethylene terephthalate (PET) (Auta et al. 2017), which are abundant in benthic environments (Wright et al., 2013). While our analyses were not designed to assess the costs of this high microplastic burden, future work in this area may help to identify the true harms of ingested plastics on cnidarian physiology.

Mixotrophy may play a crucial role in the resilience of jellyfish in the genus *Cassiopea*. Larson (1997) highlighted the importance of heterotrophic feeding in *Cassiopea*, observing ingestion of small epibenthic crustaceans. The contributions of endosymbiotic dinoflagellates (Family Symbiodiniaceae) to *Cassiopea*’s nutrition should not be underestimated, as symbiont-derived carbon can meet a significant portion of the host’s requirements (Lyndby et al., 2020; Toullec et al., 2024). The low gut content observed in Floridian bleached jellyfishes in this study suggest heterotrophy may not confer as much resilience after bleaching. Identifying whether heterotrophic capacity suffers at high temperatures within *Cassiopea* will determine the resiliency to be expected of these medusae.

## 5. CONCLUSION

In this study, we analyzed the gut content of the benthic jellyfish *Cassiopea* from 8 locations worldwide. Our results support the hypothesis that shallow water *Cassiopea* from across the globe have some shared food items and has a strong benthic component. Moreover, our data could be used as baseline to generate reasonable estimates for wet mass associated with *Cassiopea* medusae. Our field data indicate that medusae may be clearing upwards of 150 small adult crustaceans daily from the water column and benthos surrounding them. *Cassiopea* represent significant contributors to local turbidity and are highly abundant across subtropical coastlines; thus, assessing basic features of consumption and prey diversity are core to positioning them within networks and food webs. This paper contributes ecological data about the trophic position of *Cassiopea,* which may support further research on their ecological impact on the environment and may indicate avenue for improving or modifying husbandry practices.

## ACKNOWLEDGEMENTS

We thank the Smithsonian Tropical Research Institute for organizing the Cnidaria taxonomy course held in Panama in 2022 through NSF OISE-1828949 (to RC), and NSF BIO 2153775 (to MPM) grants, which allowed us to collect the first set of data on gut content. We thank the organizers of the Jellyfish Bloom Symposium (JB7), thanks to which we were able to find more researchers interested in this topic and willing to collaborate. We also thank the organizers of the international *Cassiopea* workshop for providing the location for collection at KLML. Kaden Muffett and Marta Mammone want to thank all the coauthors for the great collaboration efforts from all over the world. Andrea Toso and Lara Fumarola want to thank the National Recovery and Resilience Plan (NRRP), Mission 4 Component 2 Investment 1.4 - Call for tender No. 3138 of 16 December 2021, rectified by Decree n.3175 of 18 December 2021 of Italian Ministry of University and Research funded by the European Union – NextGenerationEU. Project code CN_00000033, Concession Decree No. 1034 of 17 June 2022 adopted by the Italian Ministry of University and Research, CUP D33C22000960007, Project title “National Biodiversity Future Center - NBFC”) and the project ACTNOW “Advancing understanding of cumulative impacts on European marine biodiversity, ecosystem functions and services for human wellbeing”, Horizon Europe, G.A. No. 101060072 for the funding of the Italian sampling; Dr. Mar Bosch-Belmar for sampling support and Dr. Emanuele Mancini for help during crustacean identification.

## 7. AUTHOR CONTRIBUTIONS

Kaden Muffett, Marta Mammone: writing original draft, supervision; Kaden Muffett, Ramón D. Morejón-Arrojo: data analysis. Kaden Muffett, Marta Mammone, Anabelle Marques Klovrza, Valentina Cardona: conceptualization, data collection. Ramón D. Morejón-Arrojo, Andrea Toso, Lara M. Fumarola, Serafin Mendez Geson III, Claire E. Rowe, Anthony Bellantuono: data collection; Maria Pia Miglietta: funding; Rachel Collin: funding, resources. All authors: review and editing.

## 8. STATEMENTS AND DECLARATIONS

**Conflicts of interests** The authors do not have any competing financial or non-financial interests to disclose. **Funding** MM wants to thank the Smithsonian Tropical Research Institute for a fellowship. **Ethics approval** All applicable international, national, and/or institutional guidelines for the care and use of animals were followed. **Data availability** The datasets generated during and/or analyzed during the current study are available from the corresponding authors on reasonable request.

