## Supplementary material for "Notes on the diet and size of *Cassiopea*": S1-3 figs

Supplementary Materials

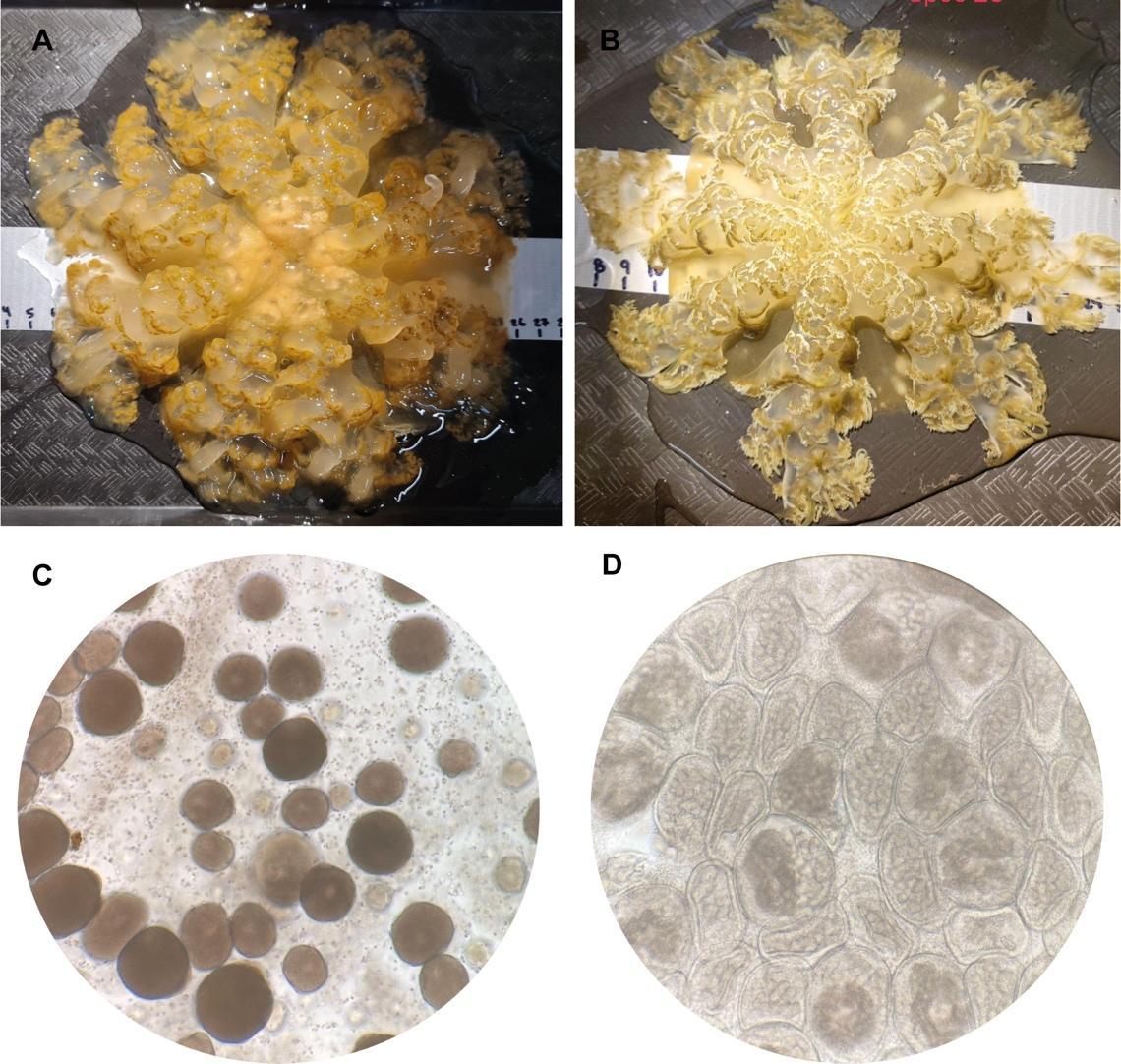

Figure S1. Female (A) and male (B) *Cassiopea* with eggs (C) and spermatocytes (D) from gonad tissue.

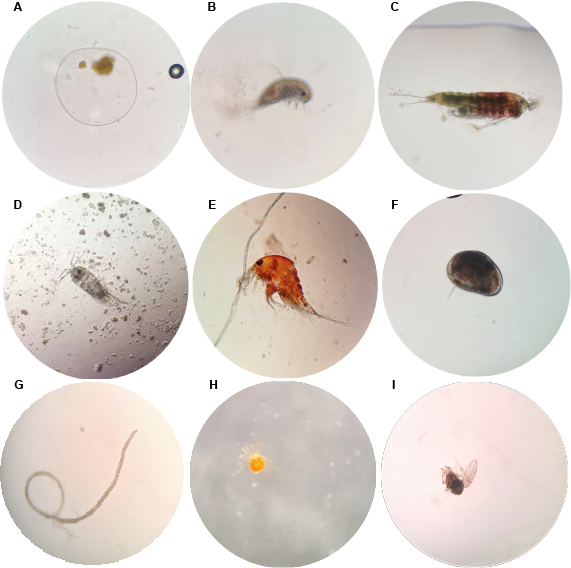

Figure S2. Examples of *Cassiopea* sp. food items. A) egg; B) amphipod; C-E) copepods; F) ostracod; G) nematode; H) water mite; I) insect.

S3: Mass Models

Power series using only adult animals

WW=0.111*(BD^2.83)
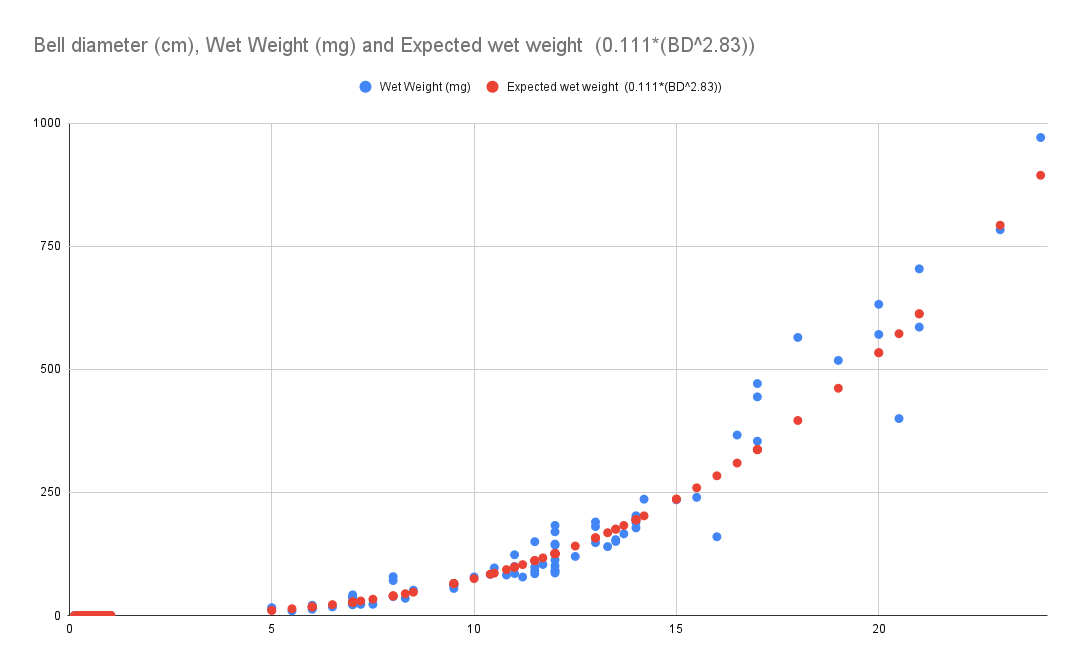

Fig. 1: Expected wet weight and actual weight by power series

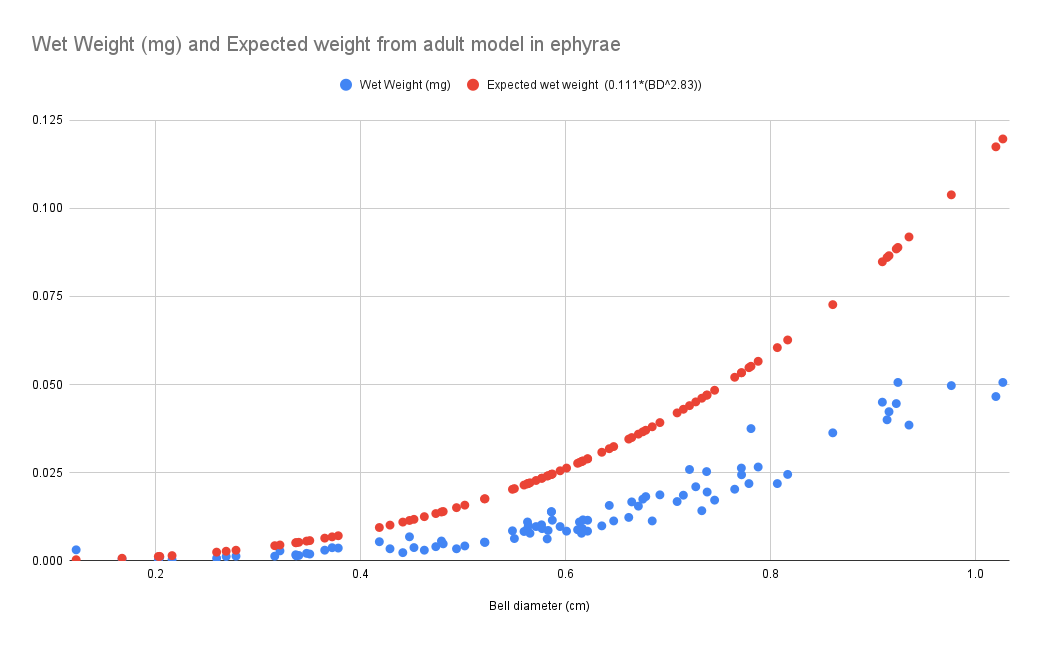

Fig. 2: Fig. 1: Expected wet weight and actual weight by power series of ephyrae

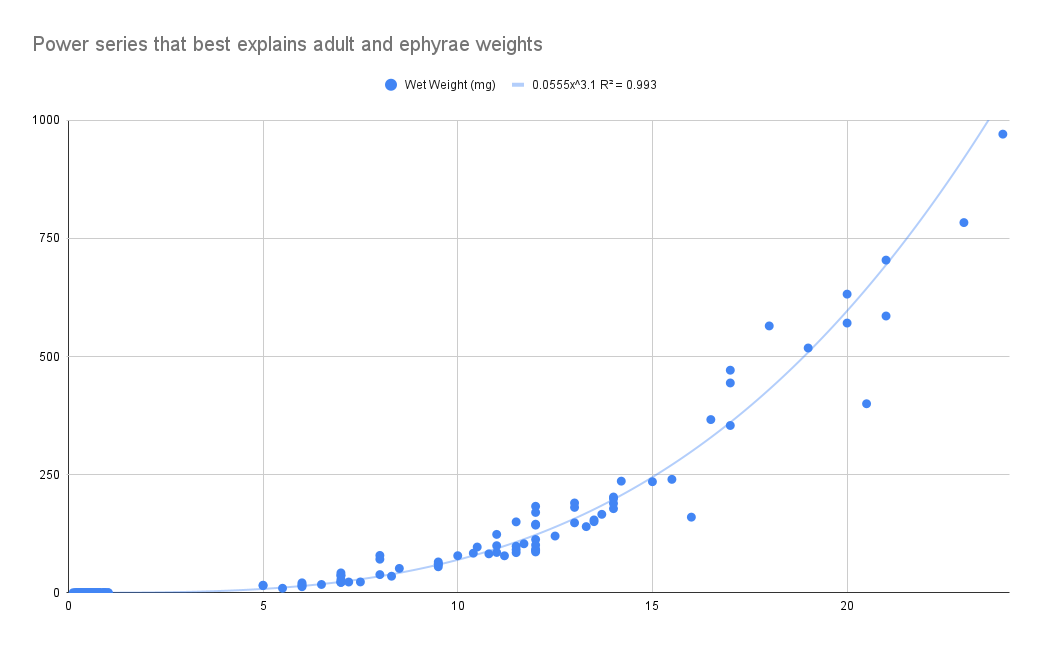

Fig 3: Actual weight and power series based onindividuals from 0~30cm
